# Absence of Lipopolysccharide (LPS) expression in Breast Cancer Cells

**DOI:** 10.1101/2023.08.28.555057

**Authors:** Noel FCC de Miranda, Vincent THBM Smit, Manon van der Ploeg, Jelle Wesseling, Jacques Neefjes

## Abstract

The association between bacterial activity and tumorigenesis has gained attention in recent years, alongside the well-established link between viruses and cancer. A recent study proposed the presence of intracellular bacteria in cancer cells, particularly in melanomas and breast cancers, with detectable bacterial DNA. The authors suggested that these bacteria contribute to the tumors’ development. We sought to replicate these findings using the same experimental methods on different tissue microarrays. Our investigation included 129 breast cancer samples, but we found no evidence of LPS expression within cancer cells. Instead, LPS immunoreactivity was observed in ducts or immune cells, specifically macrophages. The discrepancies in LPS staining warrant caution in interpreting the reported observations, and further research is needed to elucidate the potential role of intracellular bacteria in cancer development.

## Brief communication

While the etiological role of viruses in cancer has long been appreciated, bringing to light associations between bacterial activity and tumorigenesis has proven more challenging. Nevertheless, groundbreaking epidemiological and cell biological studies have, in recent years, connected particular bacterial pathogens such as *Salmonella Typhi* and *Helicobacter pylori* to gallbladder carcinoma and gastric cancer, respectively (*1, 2*). Furthermore, bacteria-derived genotoxic signatures have been observed in colorectal cancer (*3, 4*).

Recently, a report proposed that many cancers presented specific intracellular bacteria in the cancer cells themselves (*5*). Bacterial ribosomal 16S gene sequencing from hundreds of different tumor tissues revealed that 14% of melanomas and 63% of breast cancers showed detectable bacterial DNA. To determine the localization of bacteria, the authors performed immunohistochemical detection of tissues with an anti-LPS antibody. LPS was detected in both cancer and immune cells. Also, 16S RNA probing was performed. These experiments suggested that most (breast) cancer cells within a tumor displayed LPS expression, interpreted as derived from intracellular bacteria.

This observation was striking as LPS is expected to induce a robust immune response, which would result in a strong inflammatory response or even a sepsis (*6*). Yet, while many breast cancer tissues in the Nejman study (*5*) strongly stained for LPS, breast cancer is not considered to be a particularly immunogenic tumor type (*7*). Secondly, the subcellular localization of LPS is striking: the images presented by the authors suggest a nuclear localization of LPS – an unprecedented location for intracellular bacteria.

These issues inspired us to test the observations reported by Nejman *et al*. (*5*). Therefore, we assessed LPS expression using the same antibody clone used in the Nejman study (WN1 222-5, HycultBiotech) on a commercial tissue microarray (BR10010f-BX, US Biomax) containing 50 cases of breast carcinoma and matched (lymph node) metastatic lesions, a tissue microarray composed at the Netherlands Cancer Institute containing 42 ductal and 15 lobular breast carcinomas, and on 22 biopsies from invasive breast cancer (at the Leiden University Medical Center). LPS immunoreactivity was discerned in 12%, 5%, and 14% of the 129 breast cancer samples, respectively. Strikingly, LPS expression was never detected in cancer cells but rather inside ducts or in immune cells identified as macrophages (Figure 1). This localization does fit with the expected pattern of LPS expression in tissues.

**Figure 1.**
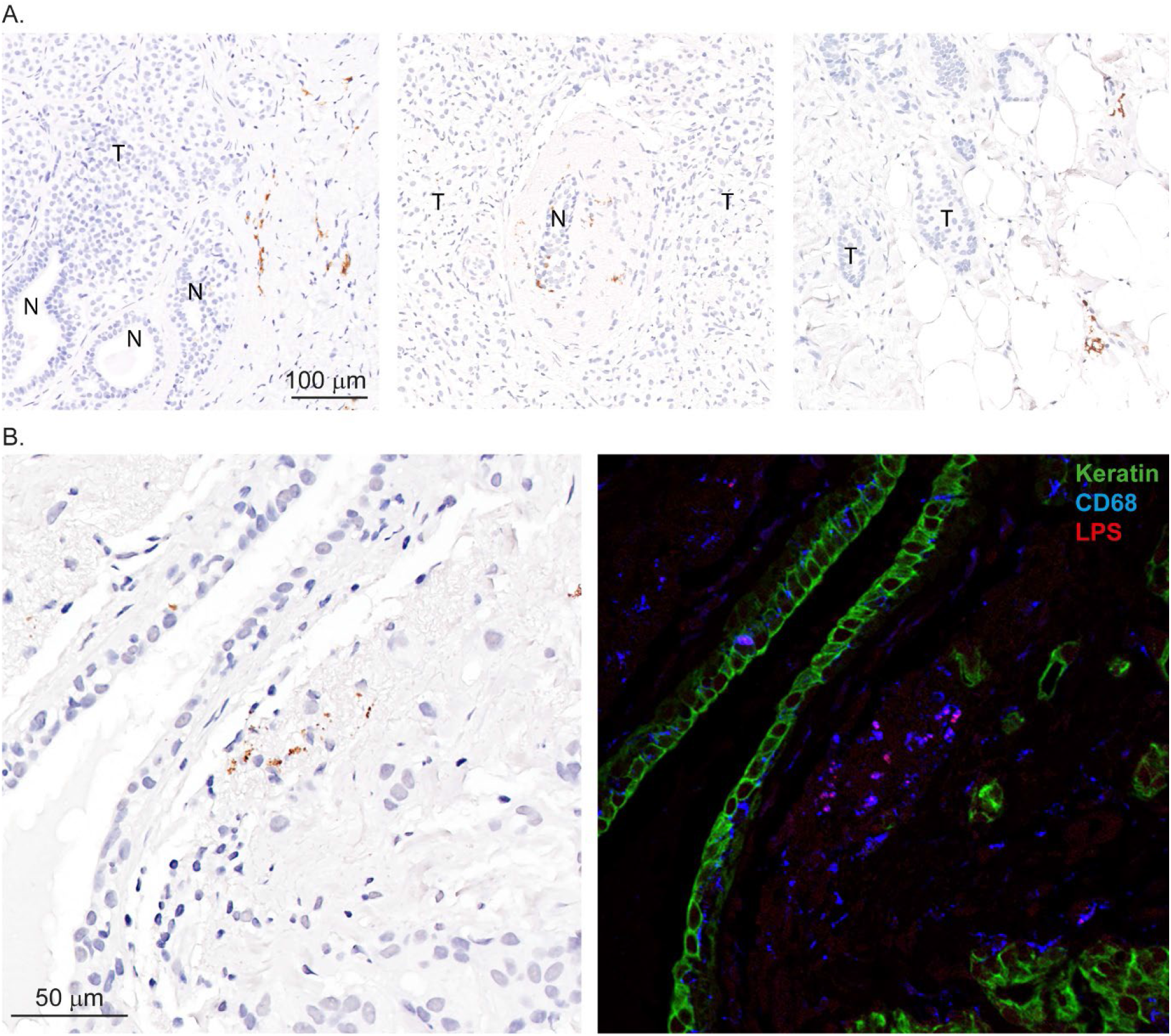
A - Representative examples of LPS immunodetection with typical granular pattern. LPS expression never co-localized with cancer cells. B – Left: LPS detection of a breast cancer section by classical anti-LPS IHC. Right panel, triple immunofluorescence of the next slice in this breast cancer tissue stained for anti-Keratin (green), anti-CD68 (blue) and anti-LPS (red) antibodies to show that LPS localized in CD68 positive cells. CD68 marks macrophages, supporting the detection of LPS inside macrophages. Bar indicates magnification.

Previously, attempts to detect LPS in gallbladder carcinomas, strongly linked to chronic *Salmonella Typhi* infections, already showed absence of LPS expression, despite detection of *S. Typhi*-derived DNA in the same samples (*1*). Therefore, we do not dispute the presence of bacterial DNA in the tumor samples, as reported by Nejman and colleagues. We do however question the immunohistochemical results upon which their central thesis rests, namely that cancer cells are ‘composed of tumor type-specific bacteria’ (*5*). The LPS stainings reported by Nejman and colleagues, claiming abundant presence of LPS in human cancer tissues and, especially, in the nucleus of cancer cells, could not be reproduced in any of the 129 breast cancer tissues analyzed. They may be the result of unspecific antibody binding and/or LPS contamination of the tissues.

Our challenge in confirming the presence of LPS in breast cancer cells adds to the current discourse on the effectiveness of microbiome-focused approaches that offer proof of bacterial components within human cancers. Specifically, recent bioinformatic investigations into genomic data identifying microbial sequences in human cancers (*8-10*) have faced criticism due to several methodological flaws and misinterpretation of findings (*11, 12*). This discussion emphasizes the importance of prudently employing bioinformatic analyses, ideally in conjunction with independent biological experiments— though such experiments should themselves be subject to meticulous scrutiny, as demonstrated by the lack of reproducibility in detecting LPS in cancer cells.

## Methods

### Patient material

We obtained three different cohorts for our study. A tissue array, BR10010f-BX, was purchased from US Biomax and comprised 50 cases of breast carcinoma along with matched (lymph node) metastatic lesions. The second tissue microarray was obtained from the Netherlands Cancer Institute and included 42 ductal and 15 lobular breast carcinomas. Additionally, we investigated 22 biopsies from invasive breast cancers diagnosed at the Leiden University Medical Center. All retrospective medical data/biospecimen studies at the Netherlands Cancer Institute have been executed pursuant to Dutch legislation and international standards. Prior to 25 May 2018, national legislation on data protection applied, as well as the International Guideline on Good Clinical Practice. From 25 May forward we also adhere to the GDPR. Within this framework, patients are informed and have always had the opportunity to object or actively consent to the (continued) use of their personal data & biospecimens in research. Hence, the procedures comply both with (inter-) national legislative and ethical standards. All samples were anonymized and handled according to the ethical guidelines described in the Code for Proper Secondary Use of Human Tissue in the Netherlands of the Dutch Federation of Medical Scientific Societies.

### Immunohistochemistry

FFPE tissue sections of 4-μm thick were deparaffinized and rehydrated followed by heat-mediated antigen retrieval using citrate buffer, pH 6.0. Slides were blocked using SuperBlock (PBS) Blocking Buffer (Thermo Scientific, Cat# 37515) for 30 minutes at RT. The anti-LPS antibody (clone WN1 222-5, HycultBiotech) was diluted 1:000 in 1% BSA/PBS solution. Tissue sections were incubated overnight at 4°C. Subsequently, slides were incubated with poly-HRP (Thermo Scientific, Cat# 21140) for 60 minutes and developed with DAB chromogen (1:50, DAKO #K3468). Sections were counterstained with hematoxylin (DiaPath #C0305). Slides were dehydrated and mounted with Micromount (Leica #3801731). The immunohistochemistry procedure was evaluated by expert pathologists, and the scoring was determined based on the presence or absence of LPS immunodetection.

## Acknowledgments

NFCCdM is supported by the European Research Council (ERC) under the European Union’s Horizon 2020 Research and Innovation Programme (grant agreement no. 852832). JN is supported by grants from the Dutch Cancer Society KWF. We would like to acknowledge the NKI-AVL Core Facility Molecular Pathology & Biobanking (CFMPB) for supplying NKI-AVL Biobank material and /or lab support.

## Notes

### Competing Interest Statement

The authors have declared no competing interest.

## References

1. T. Scanu et al., Salmonella Manipulation of Host Signaling Pathways Provokes Cellular Transformation Associated with Gallbladder Carcinoma. Cell Host Microbe 17, 763–774 (2015).

2. L. E. Wroblewski, R. M. Peek, Jr., K. T. Wilson, Helicobacter pylori and gastric cancer: factors that modulate disease risk. Clin Microbiol Rev 23, 713–739 (2010).

3. C. Pleguezuelos-Manzano et al., Mutational signature in colorectal cancer caused by genotoxic pks(+) E. coli. Nature 580, 269–273 (2020).

4. P. J. Dziubanska-Kusibab et al., Colibactin DNA-damage signature indicates mutational impact in colorectal cancer. Nat Med 26, 1063–1069 (2020).

5. D. Nejman et al., The human tumor microbiome is composed of tumor type-specific intracellular bacteria. Science 368, 973–980 (2020).

6. C. V. Rosadini, J. C. Kagan, Early innate immune responses to bacterial LPS. Curr Opin Immunol 44, 14–19 (2017).

7. J. P. Bates, R. Derakhshandeh, L. Jones, T. J. Webb, Mechanisms of immune evasion in breast cancer. BMC Cancer 18, 556 (2018).

8. L. Narunsky-Haziza et al., Pan-cancer analyses reveal cancer-type-specific fungal ecologies and bacteriome interactions. Cell 185, 3789–3806 e3717 (2022).

9. G. D. Poore et al., Microbiome analyses of blood and tissues suggest cancer diagnostic approach. Nature 579, 567–574 (2020).

10. B. Chrisman et al., The human “contaminome”: bacterial, viral, and computational contamination in whole genome sequences from 1000 families. Sci Rep 12, 9863 (2022).

11. A. Gihawi, C. S. Cooper, D. S. Brewer, Caution regarding the specificities of pan-cancer microbial structure. Microb Genom 9, (2023).

12. A. Gihawi et al., Major data analysis errors invalidate cancer microbiome findings. bioRxiv, (2023).

